# Everything you wanted to know about Mayaro virus but were afraid to ask: Characterization and lifecycle of Mayaro virus in vertebrate and invertebrate cellular backgrounds

**DOI:** 10.1101/2020.11.18.388884

**Authors:** Sujit Pujhari, Marco Brustolin, Chan C. Heu, Ronald Smithwick, Mireia Larrosa, Susan Hafenstein, Jason L. Rasgon

**Affiliations:** Department of Entomology, Center for Infectious Disease Dynamics and the Huck Institutes of the Life Sciences, The Pennsylvania State University, University Park, PA; Department of Pharmacology Physiology and Neuroscience, University of South Carolina School of Medicine, Columbia, South Carolina, USA; harmacology, Physiology, and Neuroscience; USDA-ARS, Maricopa, AZ, USA; Universitat Autònoma de Barcelona, Spain; Department of Biochemistry and Molecular Biology,, The Pennsylvania State University, University Park, PA; Department of Medicine, The Pennsylvania State University College of Medicine, Hershey, PA, USA

**Author notes:** Address of correspondence to Jason L. Rasgon and Sujit Pujhari.

**Keywords:** Arbovirus, Mayaro virus, macropinocytosis, cytopathic vacuoles, virus-host interactions

## Abstract

Mayaro virus (MAYV) is an emerging new world alphavirus (genus *Alphavirus*, family *Togaviridae)* that causes acute multiphasic febrile illness, skin rash, polyarthritis, and occasional severe clinical phenotypes. The virus lifecycle alternates between invertebrate and vertebrate hosts. Here we characterize the replication features, cell entry, life cycle, and virus-related cell pathology of MAYV using vertebrate and invertebrate *in vitro* models. Electron dense clathrin-coated pits in infected cells, and reduced viral production in the presence of dynasore, ammonium chloride, and bafilomycin, indicates that viral entry occurs through pH-dependent endocytosis. Increase in FITC-dextran uptake (an indicator of macropinocytosis) in MAYV-infected cells, and dose-dependent infection inhibition by 5-(N-ethyl-N-isopropyl) amiloride (a macropinocytosis inhibitor), indicated that macropinocytosis is an additional entry mechanism of MAYV in vertebrate cells. Acutely infected vertebrate and invertebrate cells formed cytoplasmic or membrane-associated extracytoplasmic replication complexes. Mosquito cells showed modified hybrid cytoplasmic vesicles that supported virus replication, nucleocapsid production, and maturation. Mature virus particles were released from cells by both exocytosis and budding from the cell membrane. MAYV replication was cytopathic and associated with induction of apoptosis by the intrinsic pathway, and later by the extrinsic pathway in infected vertebrate cells. Given that MAYV is expanding its geographical existence as a potential public health problem, this study lays the foundation of biological understanding valuable for therapeutic and preventive interventions.

## Introduction

Mayaro virus (MAYV) is a neglected emerging arboviral pathogen. It was first isolated in 1954 from Trinidad and Tobago, and since then, outbreaks have been reported in South and Central America [1]. Many of its clinical features, including arthralgia, overlap with Dengue and Chikungunya; however, biphasic or intermittent hyperthermia can distinguish MAYV from other arboviral infections [2, 3]. MAYV can cause neurological complications, myocarditis, hemorrhagic manifestations, and death [2, 4]. MAYV alternates between vertebrate and invertebrate hosts and is primarily transmitted through the bite of female *Haemagogus* (in sylvatic cycle) and *Aedes* (urban and peri-urban cycle) mosquito species in South and Central America [5-7]. Transmission by multiple *Anopheline* mosquito species has also been demonstrated through laboratory studies, indicating a potential risk of this emerging virus in other parts of the world [8] (Fig.1).

**Figure 1.**
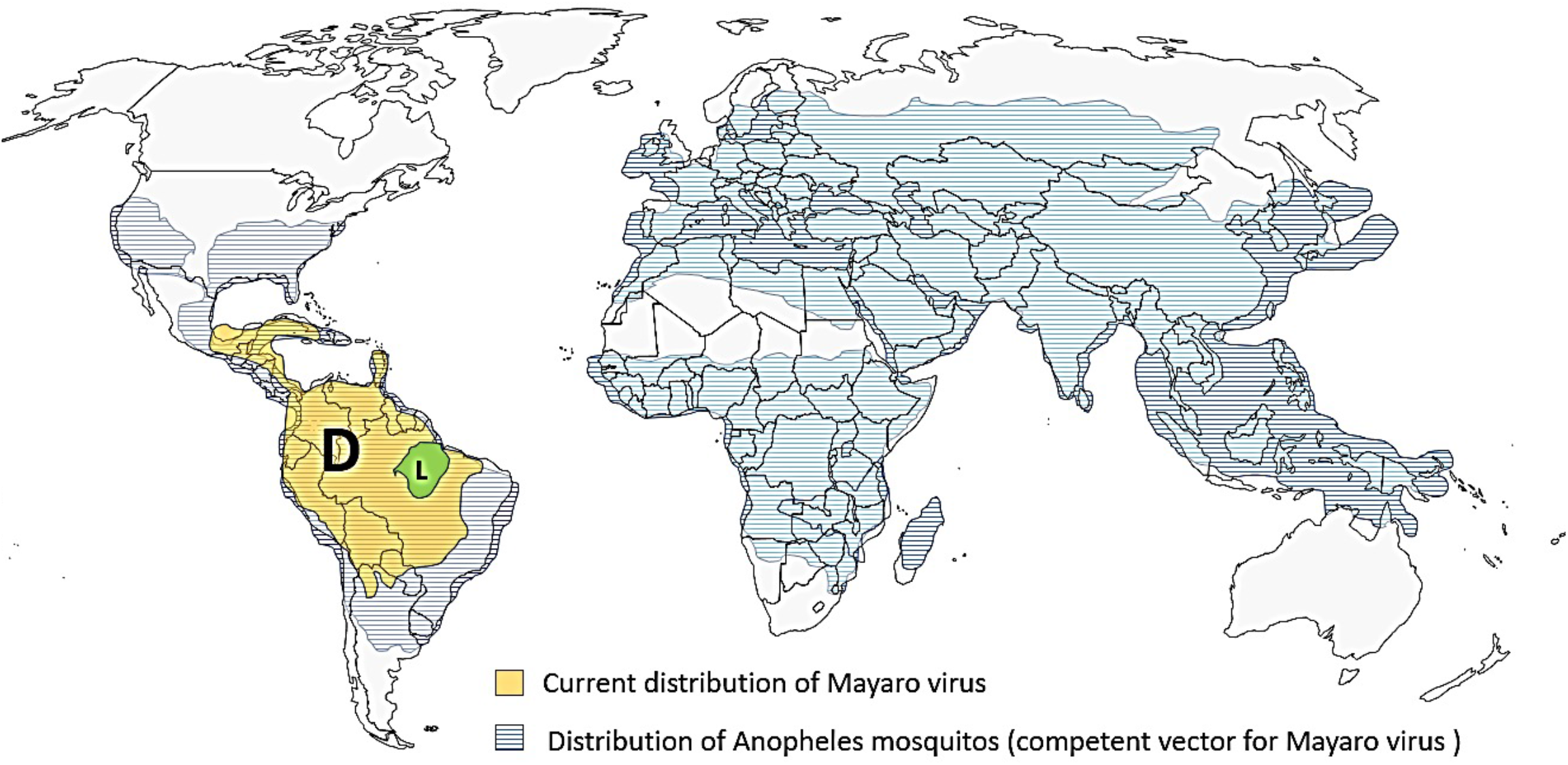
Distribution of Mayaro virus in the America and global distribution of *Anopheles* vectors. Distribution of D (yellow) and L (green) strains of Mayaro virus in South America and geographical areas at risk of future MAYV outbreaks highlighted in blue.

MAYV is a positive-sense, single-stranded-RNA virus that belongs to the genus *Alphavirus* in the family *Togaviridae*. It is a member of the Semliki Forest virus antigenic complex that consists of eight other viruses: Semliki Forest, Chikungunya (CHIKV), Bebaru, Getah, Ross River (RRV), O’nyong-nyong (ONNV), Sagiyama and Una viruses [9]. Its genome is approximately 11.7 kb and encodes four nonstructural proteins (nsP1-4), six structural proteins (capsid [C], envelope [E] proteins [E3, E2, E1], 6K, and transframe), and two open reading frames (ORFs) [10]. Based on its whole-genome phylogeny, MAYV has three genotypes (D, L, and N) which are highly conserved with approximately 17% nucleotide divergence across all three genotypes, and 4% among D strains. It is thought that genotypes D and L diverged approximately 150 years ago, and genotype N diverged approximately 250 years ago [11]. Genotype D has a diverse distribution in South America and the Caribbean, genotype L was detected in certain parts of Brazil whereas N genotype was found only in a localized region in Peru [12]. The ability of MAYV to recombine with other strains and related viruses may arise in new lineages [32].

Most of our understanding of the cellular and molecular biology of MAYV is based on studies with other alphaviruses. To bridge this critical knowledge gap on the biology of this emerging yet neglected arthritis-causing alphavirus, we used mosquito and vertebrate cells to characterize its biology. This study provides a comprehensive investigation on the lifecycle of MAYV, its replication characteristics, and cellular tropism to provide insight into the interaction of MAYV with its mosquito and vertebrate host. Given that MAYV is expanding its geographical existence as a potential public health problem, this study will lay down the foundation of biological understanding valuable for therapeutic and preventive interventions.

## Materials and Methods

### Cell culture

C6/36 (*Aedes albopictus*) cells, Aag2 (*Aedes aegypti*), and Sua5b (*Anopheles gambiae*) cells were maintained in Schneider′s insect cell culture medium. Vero (African green monkey kidney), BHK-21 (Baby Hamster Kidney), and Huh7.5 (Human liver) cells were maintained in Dulbecco’s modified Eagle’s medium (DMEM) supplemented with 10% fetal bovine serum (FBS), 50 units ml^-1^ of penicillin and 50 μg ml^-1^ of streptomycin. Vertebrate cells were cultured in a 37 °C incubator with 5% CO_2_, while invertebrate (mosquito) cells were cultured at 28 °C incubator without CO_2_.

### Antibodies

CHIK-48, anti-E2 protein (BEI resources, USA); anti-dsRNA, dsRNAJ2 (SCICONS, Hungary); Cleaved caspase-3, 9664, (Cell Signaling Technology [CST], USA); Cleaved Caspase-9, 52873 (CST, USA); Caspase-8, SAB3500404 (Sigma, USA); PARP, 9542 (CST, USA); HSP60, SAB4501464 (Sigma, USA); Phalloidin-594; actin (Invitrogen, USA).

### Viral growth kinetics by focus forming assay (FFA)

Approximately 3 × 10^4^ Vero cells per well were seeded in a 96-well plate and incubated overnight. Virus samples (BeAn 343102, D strain of MAYV) were diluted in ten-fold serial dilutions in DMEM without FBS supplemented with antibiotics; 30 μl from each dilution was added to each well containing the cells and incubated at 37°C. One hour post infection, 100 μl of overlay media (1X DMEM medium, 10% FBS, 50 units ml–1 penicillin, 50 μg ml–1 streptomycin, 1% carboxymethyl cellulose) was added. Twenty-four hours post-infection, the overlay was removed; to fix the cells, 100 μl of 4% paraformaldehyde in phosphate buffered saline (PBS) was added and incubated at RT for 15 minutes. For antibody probing, the plate was first blocked for 30 minutes at RT with 50 μl of blocking solution (3% BSA, 0.25% triton x in PBS), followed by the addition of 30 μl of primary antibody (CHIK-48, anti-E2 protein) for 2 h at RT or overnight at 4°C. Plates were then washed three times with PBS. Secondary antibody (Alexa 488 Goat anti-Mouse, ThermoScientific) was added (30 μl per well) and incubated for 1 h at RT. Plates were washed three times with distilled water, air-dried and screened manually under 4x objective of Olympus microscope. Viral titers were expressed as FFU ml^−1^.

### Heat treatment of virus particles

Approximately 10^6^ FFU of MAYV, Sindbis virus (SINV) and O’nyong yong virus (ONNV) were subjected to a thermal gradient treatment from 30 to 60 °C for 3 h with a thermocycler (Bio-Rad T100 Thermal Cycler), after which samples were immediately titrated on Vero cells. A non-heat-treated virus control kept at 4°C for 3 h was also included. The ratio of the number of FFU in heat-treated versus the non-heat-treated viruses was calculated to determine the relative infectivity.

### Plaque assay

Vero cells (5×10^5^ cells/well) were grown overnight to a confluent monolayer in 6 well plates and infected with serial dilutions of MAYV or ONNV or SINV-infected culture supernatant. Virions were allowed to adsorb on the cell surface for 1 hour at 37°C with 5% CO_2_; subsequently, monolayers were rinsed with DMEM without FBS, and overlay medium (1% methylcellulose in DMEM with 5% FBS) was added. The plates were incubated at 37 °C in a 5% CO_2_ incubator for 72 h. At the end of the incubation period, overlay media was removed and fixed with 1 ml of 4% PFA solution at RT for 15 minutes. After one washing with PBS, methylene blue prepared in methanol was added onto the fixed cells. After 30 minutes of incubation, plates were cleaned in tap water, and plaques size were measured.

### Effect of lysosomotropic drugs on MAYV entry

BHK-21 (3×10^4^ cells/well), Huh7.5, and C6/36 (5×10^4^ cells/well) cells were seeded in 96 well plates the day before treatment. Cells were pretreated for 3h in serum-free media containing ammonium chloride or bafilomycin A1 (see Results for concentrations used). Following incubation, cells were infected with MAYV at an MOI of 1 in the presence of each compound for 1h at 37°C. Cells were washed, complete media containing each compound was added, and were incubated at 37°C for 16h. Additionally, no drug and drug with no virus control were included. After incubation, cells were fixed with 4% PFA, permeabilized, processed for immunofluorescence using virus-specific antibodies, and virus-positive cells were quantified.

### Immunofluorescence analysis of infected cells

Huh7.5 or C6/36 cells were seeded in two well chamber slides at a density of 2 × 10^5^ cells per well. Cells were then infected with MAYV at an MOI of 1 or mock-infected and incubated at 37 °C (Huh7.5) or 28 °C (C6/36). After 1 h, cells were washed with DMEM or Schneider′s insect cell culture medium without FBS and replaced with fresh growth medium and then incubated at 37 °C (Huh7.5) or 28 °C (C6/36). Cells were fixed with 4% PFA for 15 minutes at RT at 12hpi for detection of dsRNA or at 24hpi for E2 protein detection. Cells were then blocked with 500μl of blocking solution (3% BSA with 0.25% Triton-X in PBS) for 30 min. at room temperature before incubation with CHIK-48 (anti-E2) (1:500) or dsRNA antibody (1:100) diluted in blocking buffer overnight. Next day cells were washed three times with PBS/T and incubated with Alexa Fluor 488 or 595 secondary antibodies (Life Technologies) diluted in PBS/T for 1 h at RT. After incubation, cells were washed three times with PBS/T. Finally, for nuclear staining, cells were incubated with 500 ul of PBS with Hoechst stain for 2 minutes, followed by a final washing with distilled water. Cells were mounted using 1.5mm cover glass with ProLong Diamond Antifade Mountant (Life Technologies). Images were taken using a Zeiss LSM 800 confocal microscope.

### TEM of infected cells

Cell pellets were fixed in 3% glutaraldehyde in PBS for 1 h at room temperature, washed with 0.1 M cacodylate buffer, and incubated in 1% OsO4 (in 0.1 M cacodylate buffer) for 40 min at room temperature. Samples were then washed once with 0.1 M cacodylate buffer and once in 80% acetone for a further incubation overnight at 4 °C in 2% uranyl acetate/80% acetone. The following day, serial dehydration and resin infiltration steps were performed as follows: 2 × 10 min with 80% acetone, 2 × 10 min with 90% acetone, 3 × 20 min with 100% acetone, 1 × 90 min with 50% Epon/50% acetone, 1 × 90 min with 75% Epon/25% acetone and 1 × 90 min with 100% Epon. Epon was replaced by fresh 100% Epon with polymerization accelerator BDMA and embedded at 65 °C for 72 h. Resin blocks were sectioned using a DiATOME Ultra Diamond Knife on a Leica EM UC7 ultramicrotome, from which 50nm sections were obtained and mounted on EM copper grids with carbon coating. Sections were post-stained in 2% uranyl acetate in water and Reynolds’ lead citrate for 1 min each and then processed for TEM imaging using a FEI Tecnai F20 S/TEM electron microscope.

### Cell cytotoxicity and viability assay

Neutral red uptake assay was used to evaluate MAYV-induced cell cytotoxicity/death and cytotoxicity of chemicals in Huh7.5 cells. In brief, Huh7.5 cells were seeded into 96-well plates at a density of 2.5 × 10^4^ cells per well and allowed to attach for 12 h. The outer perimeter wells of the plate were left blank as they often have decreased cell growth. Cells were infected with 0.1 or 1 MOI of MAYV at different time points in order to harvest the plates at 3, 6, 12, 24, 36, and 48 h of post-infection. Cell plates were washed once and replenished with 100ul of neutral red medium (40 ug ml^-1^) which was prepared a day before and incubated at 37°C. After 2 h of incubation at 37 degree, plates were washed (with PBS) and 150 ul of neutral red destain solution (56% ethanol [96% concentration], 49% deionized water, 1% glacial acetic acid) was added to each well. To extract the neutral red from the cells, plates were agitated for 10 minutes on a microtiter plate shaker. The optical density of the plates was measured at 540 nm in a microtiter plate reader spectrophotometer. Each plate had blanks that contained no cells and cells without virus as no treatment reference. Cell cytotoxicity was measured using the following formula % viable cells = (Abs_samp_ - Abs_blank_) / (Abs_control_ – Abs_blank_) x 100.

### Western blot

MAYV infected and mock infected cells were harvested at the time points stated above by a cell scraper and pelleted by centrifugation. Cell pellets were washed twice with PBS and lysed in RIPA buffer (20 minutes on ice) with protease inhibitor cocktail. The lysate was cleared by centrifugation at 14,000 rpm for 20 min at 4°C and resolved on 10% or 12% sodium dodecyl sulfate polyacrylamide gel electrophoresis (SDS/PAGE). Proteins were transferred to nitrocellulose membranes (0.45, Bio-Rad). Membranes were blocked in 5% milk in TBS-tween 20 (50 mM Tris-HCl pH 7.4, 250 mM NaCl, 0.1% Tween-20) for 1 hour. Membranes were probed with primary antibody overnight at 4°C, then with corresponding HRP-conjugated secondary antibodies in 5% milk/TBS-tween. Signals were detected with the enhanced chemiluminescence method (GE healthcare).

### Infection Center assay

At 6hpi Huh7.5 cells were washed, suspended with the aid of trypsin, centrifuged, and resuspended in DMEM supplemented with ZVAD-FMK or DMSO. The infected cells were counted and diluted. Then 0.03 ml of the infected cell dilution was added onto the Vero cell monolayer prepared on the 96 well culture plate. To permit the infected cells to settle down on the Vero cells, the medium was removed after 2 hours of incubation, and 100ul of methylcellulose overlay supplemented with ZVAD-FMK or DMSO was added. At 24hpi, cells were fixed and processed as described for focus forming assay and pictured using an epifluorescence microscope. The diameters of the foci were measured using ImageJ with arbitrary units.

## Results

### Growth kinetics of MAYV in vertebrate and invertebrate cells

To determine the *in vitro* host range, growth kinetics, production of infectious viral particles, and cytopathology of MAYV, single- and multi-step growth curve analyses were performed in mammalian and insect cells (Fig. 2A). The low MOI (0.1) growth curve demonstrates the release of infectious viruses in two bursts. The first burst appears between 6-12 hours post-infection (hpi) in mammalian cells and after 12hpi in mosquito cells. The second burst appears approximately at 24hpi in Vero and BHK-21 cell and after 30hpi for Huh 7.5 and mosquito cells. It is also important to note that, except for BHK-21, other vertebrate and invertebrate cells showed a distinctive latent, exponential, and plateau phase.

**Figure 2.**
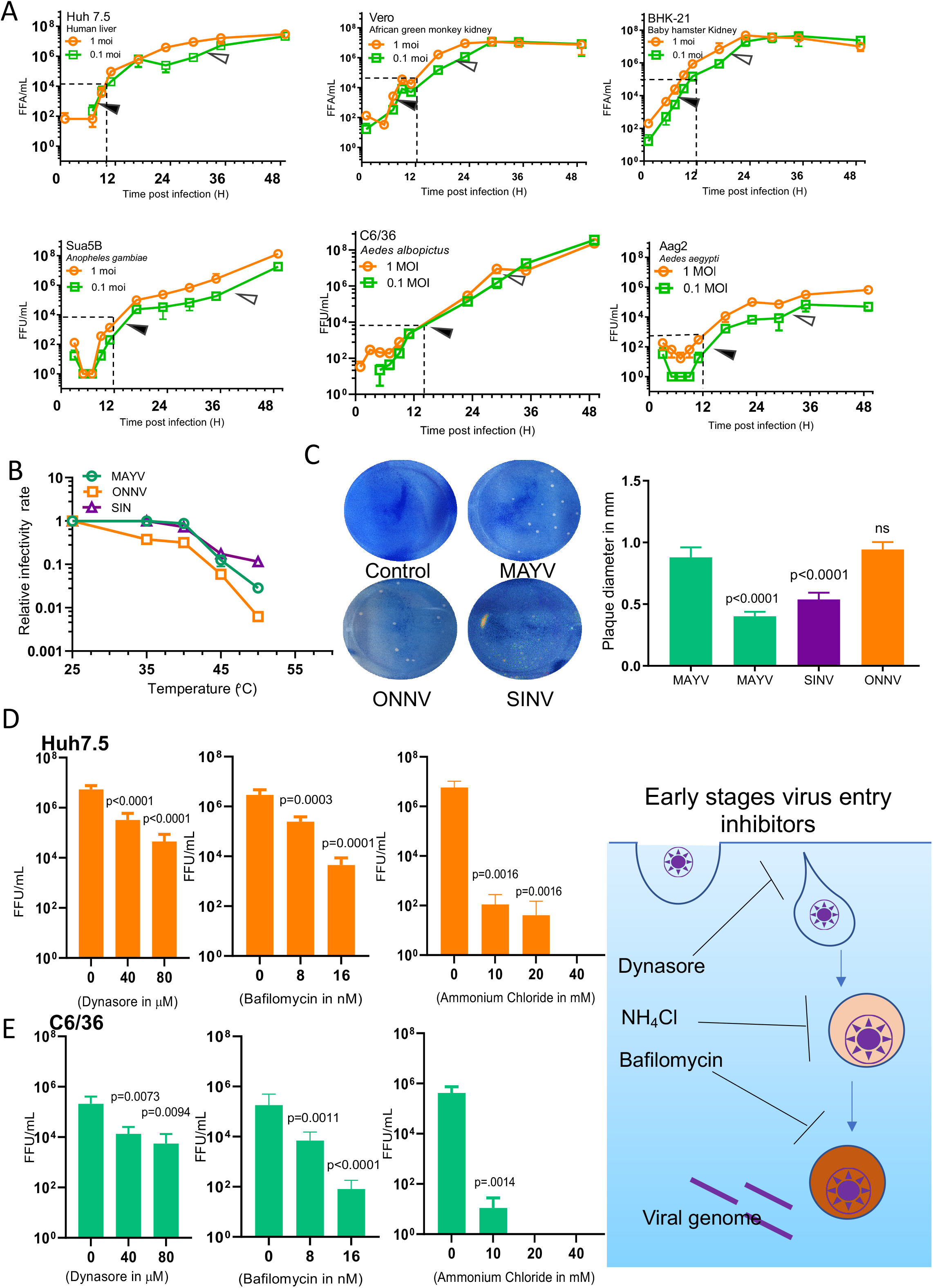
Growth kinetics, thermostability, plaque morphology and cellular entry of Mayaro virus. A) Vertebrate (Huh7.5; Human, Vero; Monkey and BHK-21; Hamster) and invertebrate (*Ae. albopictus*; C6/36, *Ae. aegypti*; Aag2 and *An. gambiae*; Sua5b) cells were infected with MAYV at an MOI of 0.1 and 1. Culture supernatants were harvested every 2-hour intervals up to 12-hour post infection, then at every 6-hour up to 36-hour post infection and a final sample was harvested at 48-hour post infection. Viral titers were quantified in the harvested samples using focus forming assay using Vero cells and growth curves were plotted. B) Mayaro, Sindbis and O’nyong’nyong virus thermostability. Stocks of indicated viruses were incubated at 4, 25, 30, 35, 40, 45, 50 and 55°C for 180 minutes and the number of infectious particles determined by focus forming assay C) Known titer stocks of MAYV, ONNV and SINV (15-20 PFU/100ul) were used to produce plaques in 6 well plates on Vero cell monolayer as detailed in the method section. The plaque diameters were measured. D and E) Effect of Dynasore, NH4Cl and Bafilomycin on MAYV entry process. D) Huh7.5 and E) C6/36 cells were treated before and after infection with the indicated doses of the chemicals. MAYV was adsorbed to cells at 1 MOI for one hour and non-adsorbed virus was removed, virus in the culture supernatant were collected at 24hpi for Huh7.5 and 36hpi for C6/36 and titrated. Data from three independent experiments were plotted, error bars indicate standard deviations of the means.

In all tested cell lines, MAYV replicated to high titers (>10^7^ FFU/mL) with the exception of the Aag2 cell line (from *Aedes aegypti*) where the maximum titer was 10^6^ FFU/ml. BHK-21 produced almost 1.5-log higher viral titer in comparison to other mammalian cells, and Sua5b produced more than 2-log higher virus titer at 24hpi and 48hpi. All mammalian cells caused cytopathic effects detectable after 12hpi which was characterized by cell rounding and detachment. Conversely, no obvious cytopathic effect was detected in mosquito cell lines.

### Physical characterization of MAYV

To understand and compare the thermal stability of MAYV to other alphaviruses such as SINV and ONNV, heat inactivation kinetics under different temperature conditions was investigated. Viral particles produced in Vero cells were treated at temperatures ranging from 25 to 55 °C for 3h, and virus infectivity was determined by focus forming assay. MAYV displayed higher thermal stability at 50°C compared to ONNV, but was lower than SINV (Fig. 2B). After 3h of incubation at 55°C all viruses were completely inactivated.

Plaque size is a measure of viral replication efficiency and genetic heterogeneity among the virus population. MAYV plaque morphology was examined in Vero cells at 72hpi and compared to ONNV and SINV plaques. MAYV formed large (∼0.8mm) and small (∼0.5 mm) plaques, while the other alphaviruses had only one type of plaque diameter (small for SINV and large for ONNV) (Fig. 2C). The large MAYV plaques were morphologically similar to ONNV, with a distinct and prominent outline. Conversely, SINV plaques were smaller, hazily outlined, and less prominent.

### pH-dependent receptor-mediated endocytic entry of MAYV

Electron dense cup-shaped structures (or pits) are prominent features during the entry phase of MAYV, as seen in electron micrographs (see below, Fig.6i, 7i). In the receptor mediated endocytosis (RME) process, these pits eventually mature and form the early endosome. At a later stage, entry features a pH-dependent fusion of the viral envelope with the endocytic vesicle and release of the viral genome into the cytoplasm.

To assess the role of RME in the entry of MAYV, we used dynasore, a pharmacological dynamin GTPase inhibitor. Dynamin is a GTPase essential for pinching off of the endosomes from the cytoplasmic membrane. Vertebrate and invertebrate cells were pretreated (1h pre-infection) with dynasore and infected with MAYV. MAYV internalization was significantly inhibited by dynasore pretreatment and its presence in a dose-dependent manner indicating the role of RME in the MAYV entry process (Fig. 2D, E).

Further, to test whether the release of MAYV into the host cytoplasm upon entry is pH-dependent, we evaluated the effect of two known pharmacological inhibitors: Ammonium chloride and Bafilomycin. The former is a weak base and a lysosomotropic agent because of its propensity to accumulate in lysosomes, and the latter blocks the v type ATPase and alters the late endosomal vesicles’ pH. Pretreatment of cells (1h pre-infection) and maintaining the chemicals in the culture medium with increasing concentrations revealed a significant dose-dependent reduction in virus particle production (Fig. 2D, E).

### Spatial distribution of viral replication complex and viral particles

To demonstrate the intracellular spatial distribution pattern of the viral replication complex, Huh-7.5 and C6/36 cells were infected with MAYV and assayed with dsRNA antibodies 6hpi. dsRNA is an intermediate in the replication complex of RNA viruses which was found to be distributed throughout the cytoplasmic compartment of both cell lines but absent in the filopodial extensions (detected using phalloidin) (Fig. 3A). Z-stack 3D analysis revealed that replication units (dsRNA puncta) were distributed both in cytoplasm and cell membrane indicated by the distribution of dsRNA puncta both in and above the plane of the nucleus (Fig. 3B, C). This corroborates with the distribution of replication vesicles both in the cytoplasm and plasma membrane (Fig. 3D), as seen under TEM (described below).

**Figure 3.**
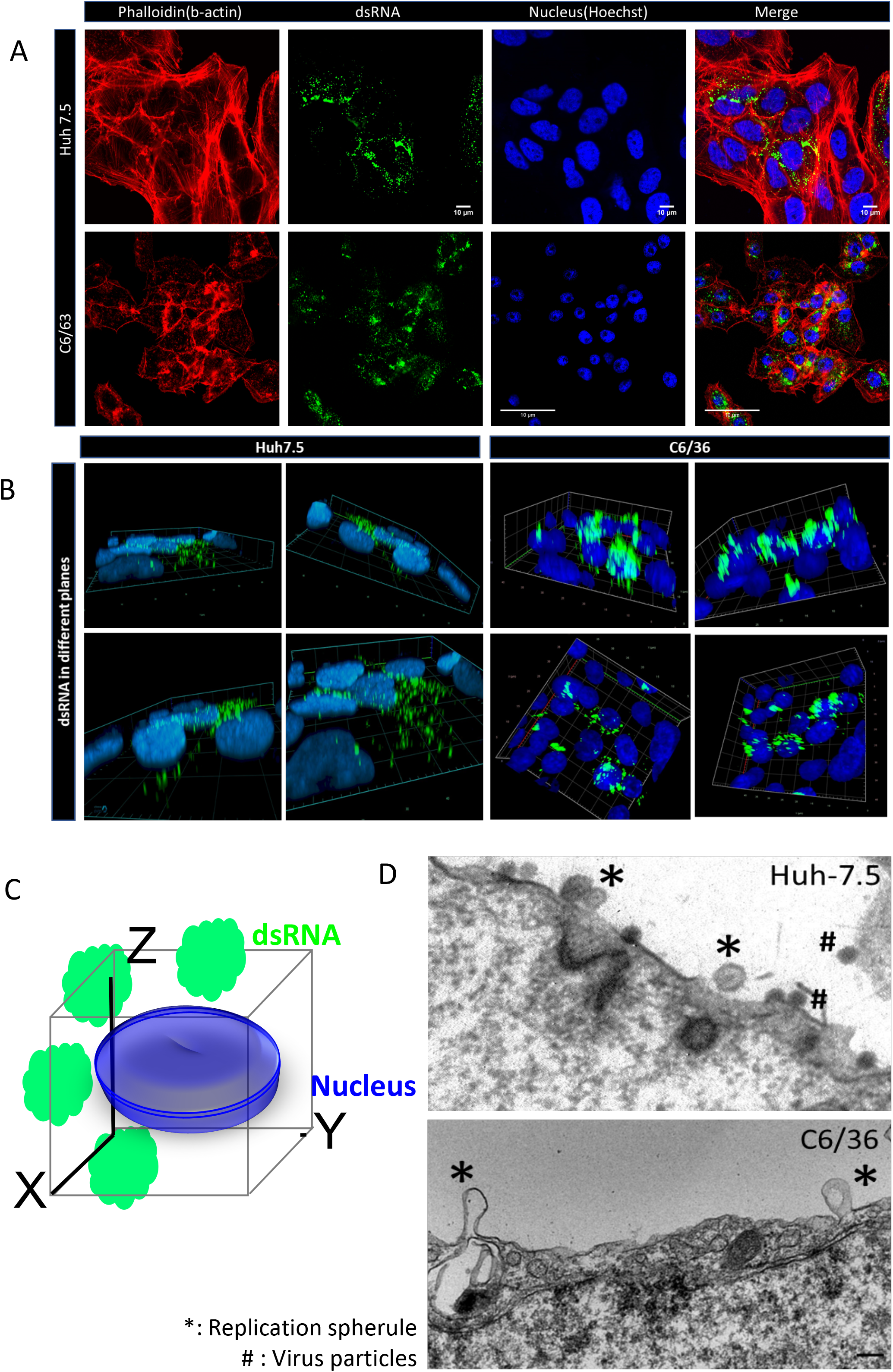
MAYV replication spherules on the plasma membrane and cytoplasmic compartment of Huh7.5 and C6/36 cells. A) Huh7.5 (upper panel) and C6/36 (lower panel) cells were infected with 1 MOI of MAYV, 6hpi cells were fixed and stained for dsRNA (green), filamentous actin (F-actin) filament with phalloidin (red) and nucleus (blue) with Hoechst stain. B) z-stack analysis of the distribution of dsRNA on diferrent planes of infected cells C) Model of expected distribution pattern of dsRNA (green), marker of the viral replication complex, at different planes in a MAYV infected cell with respect to the nucleus. D) A) Huh7.5 and C6/36 cells infected with MAYV at 5 MOI processed and scanned under TEM showing bulb like replication spherules on the cell surface.

To detect the induction of filopodial nanofiber-like extensions in MAYV infected cells and its ability to infect neighboring cells, a confocal immunofluorescence analysis was performed. Phalloidin and a CHIKV cross-reactive antibody were used to detect F-actin and MAYV E2 glycoprotein, respectively, in Huh-7.5, and C6/36 mosquito cell lines 24hpi MAYV infection. Both cell types exhibited nanofiber-like extensions connecting neighboring cells with a high density of viral particles at the surface of the membrane (Fig. 4, extreme right panel). MAYV E2 glycoproteins were primarily localized at the cell membrane and throughout the cytoplasm. Nevertheless, partial co-localization between E2 and actin was found in discrete areas (Fig. 4). This indicates that MAYV may transmit from cell-to-cell via filopodia.

**Figure 4.**
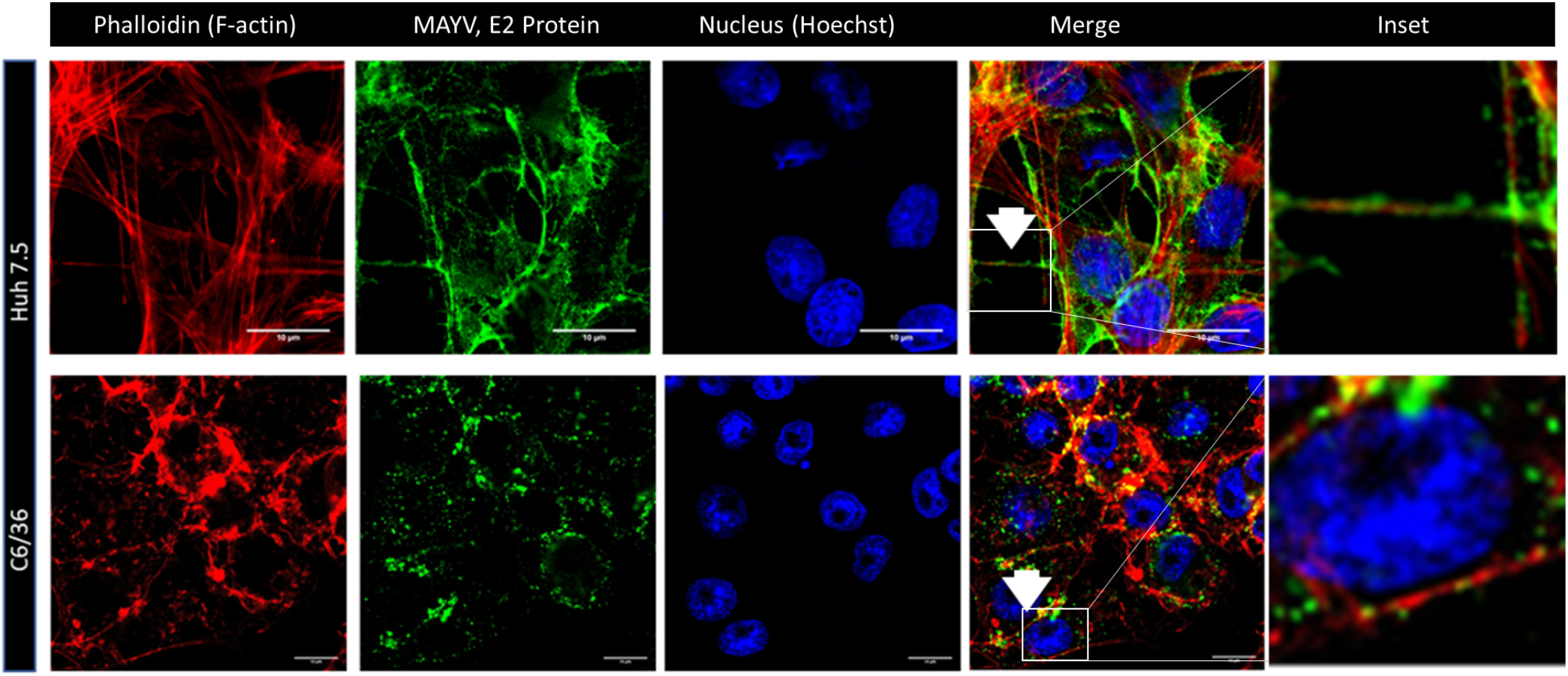
Cytoplasmic distribution of MAYV and cell-to-cell migration through nanofiber like structures in Huh7.5 and C6/36 cells. MAYV on intercellular extensions: Huh7.5 (upper panel) and C6/36 (lower panel) cells were infected with MAYV, incubated at 37°C for 12 h for Huh7.5 and 24h for C6/36 cells, and fixed. Cells were permeabilized and stained for viral E2 envelope protein (green) and phalloidin to detect F-actin (red) and nucleus (blue) with Hoechst stain. Images from one optical section are shown and are representative of three independent experiments.

### MAYV utilizes macropinocytosis for its entry

Filopodial cup-like extensions engulfing virus particles were spotted during the ultrastructural study in Huh7.5, BHK-21, and C6/36 cells, suggesting a process of macropinocytosis (Fig. 5A). To assess the involvement of macropinocytosis in MAYV entry, a functional fluid uptake assay was performed. Both vertebrate (BHK-21 and Huh-7.5) and invertebrate (C6/36 and Aag2) cell lines were incubated with 1uM FITC-dextran, and intake of FITC labeled dextran evaluated by detection of fluorescence signal in the presence or absence of MAYV. An increased FITC-dextran uptake was noted in vertebrate cells infected with MAYV, but no changes were detected in mosquito cells (Fig. 5B).

**Figure 5.**
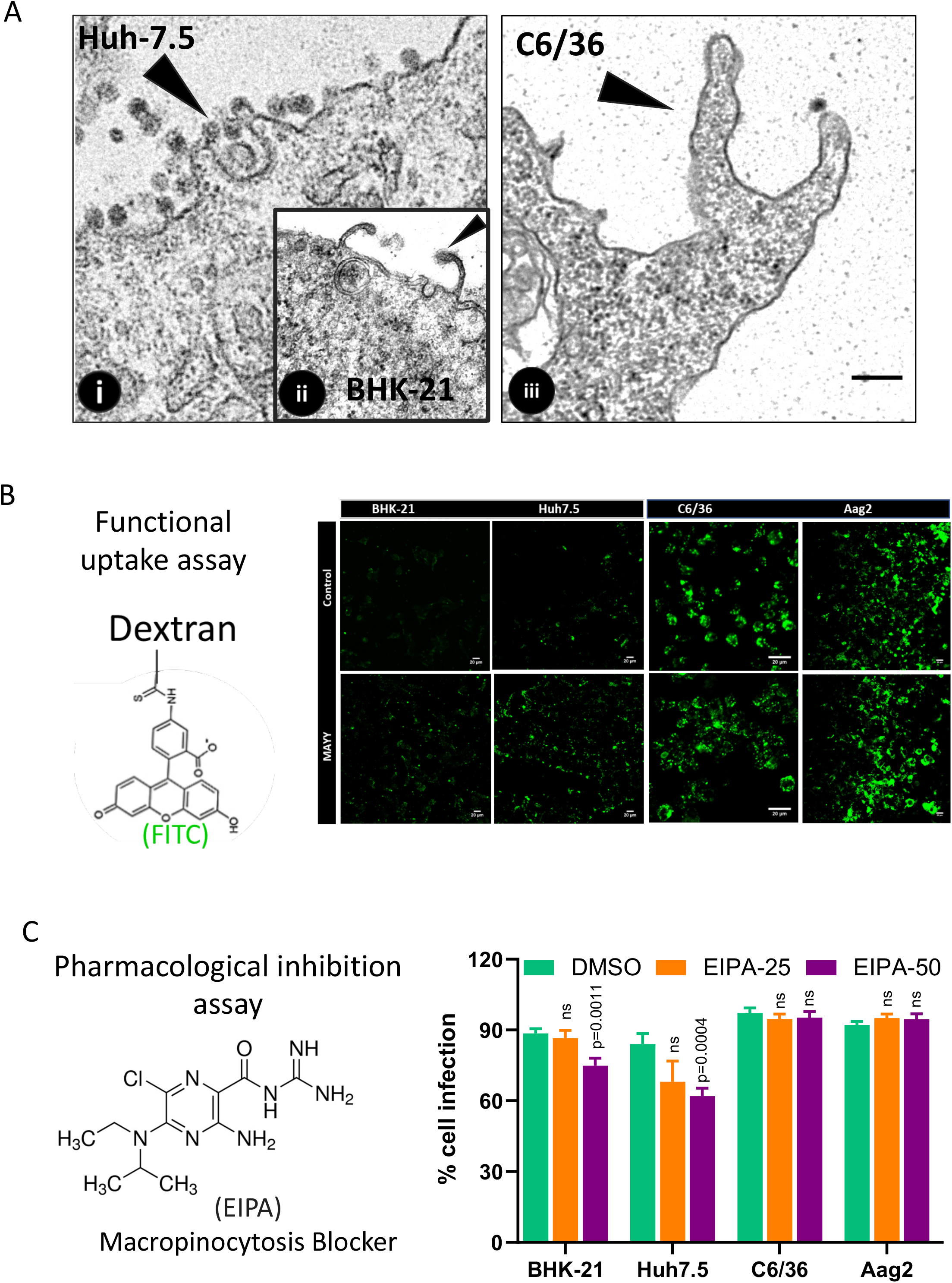
Macropinocytosis as an additional entry mechanism for MAYV in vertebrate hosts. A) Macropinocytic membrane projections chasing virus particles. Huh7.5, BHK-21 and C6/36 cells infected with MAYV at 5 MOI; 3hpi cells were harvested, processed, sectioned and scanned under TEM. B) MAYV enhances FITC-dextran uptake in vertebrate cells but not mosquito cells. BHK-21, Huh7.5, C6/36 cells were pre-treated with MAYV (lower panel) at MOI of 1 or mock infected for one hour. Cells were then washed and incubated with medium containing FITC-labeled dextran 10,000 MW (1 mg/ml). After 20 minutes, cells were washed, fixed and imaged. C) EIPA, macropinocytosis inhibitor inhibits infection of MAYV in vertebrate cells but not mosquito cells. Vertebrate and invertebrate cells were pre-treated with the different concentrations of EIPA, followed by incubation with MAYV at MOI of 0.1 in the continued presence of the drug. Control cells received DMSO instead of the drug. Cells were washed and fixed and probed with appropriate primary and secondary antibodies after 12hpi for vertebrate cells and 24hpi for invertebrate cells. Percent virus infection was calculated and plotted as bar diagram.

For further validation, a quantitative MAYV infection assay was performed in the presence of EIPA, a pharmacological inhibitor of macropinocytosis that inhibits Na+/H+ exchange. A dose-dependent reduction in infection in both BHK-21 and Huh-7.5 cells was observed, but not was not observed in mosquito cells, reinforcing the role of macropinocytosis as an important entry pathway of MAYV in vertebrate cells (Fig. 5C).

### Ultrastructural analysis of Huh7.5 and C6/36 cells infected with MAYV

To provide ultrastructural details, thin-section transmission electron microscopy of Huh7.5 and C6/36 cells infected with MAYV was performed (Fig. 6). In Huh7.5 cells, it was observed that MAYV replicated (Fig. 3D) in replication spherules, matured in the cytoplasmic vesicles (CPV) and egressed by both budding and exocytosis. The inner membrane of CPV-I, prominent in the BHK-21 MAYV infected cells, associated with bulb-shaped spherules, corresponds to the invaginations of the vacuole membranes. Inside the spherule, a central dense mass of possibly replicating viral RNA was often seen with a narrow neck connected it to the cytoplasm (Fig. 6iii). Spherules protruding toward extracellular space were also observed (Fig. 3D). Nucleocapsids (NCs) are associated with type II cytoplasmic vesicles (CPV-II) containing multivesicular inclusions, amorphous material, and occasional intact-looking virions (Fig. 6iv). Different types of CPV-II were seen coupled with NCs on the cytoplasmic side of the vacuole and/or the interior of the double-membrane CPV-II. CPV-I and CPV-II were both present in the infected cells 6hpi, though CPV-II was more abundant at 12hpi. Release of matured viral particles by budding from PM as well as through exocytosis were evident on the TEM micrographs (Fig.6 v,vi).

**Figure 6.**
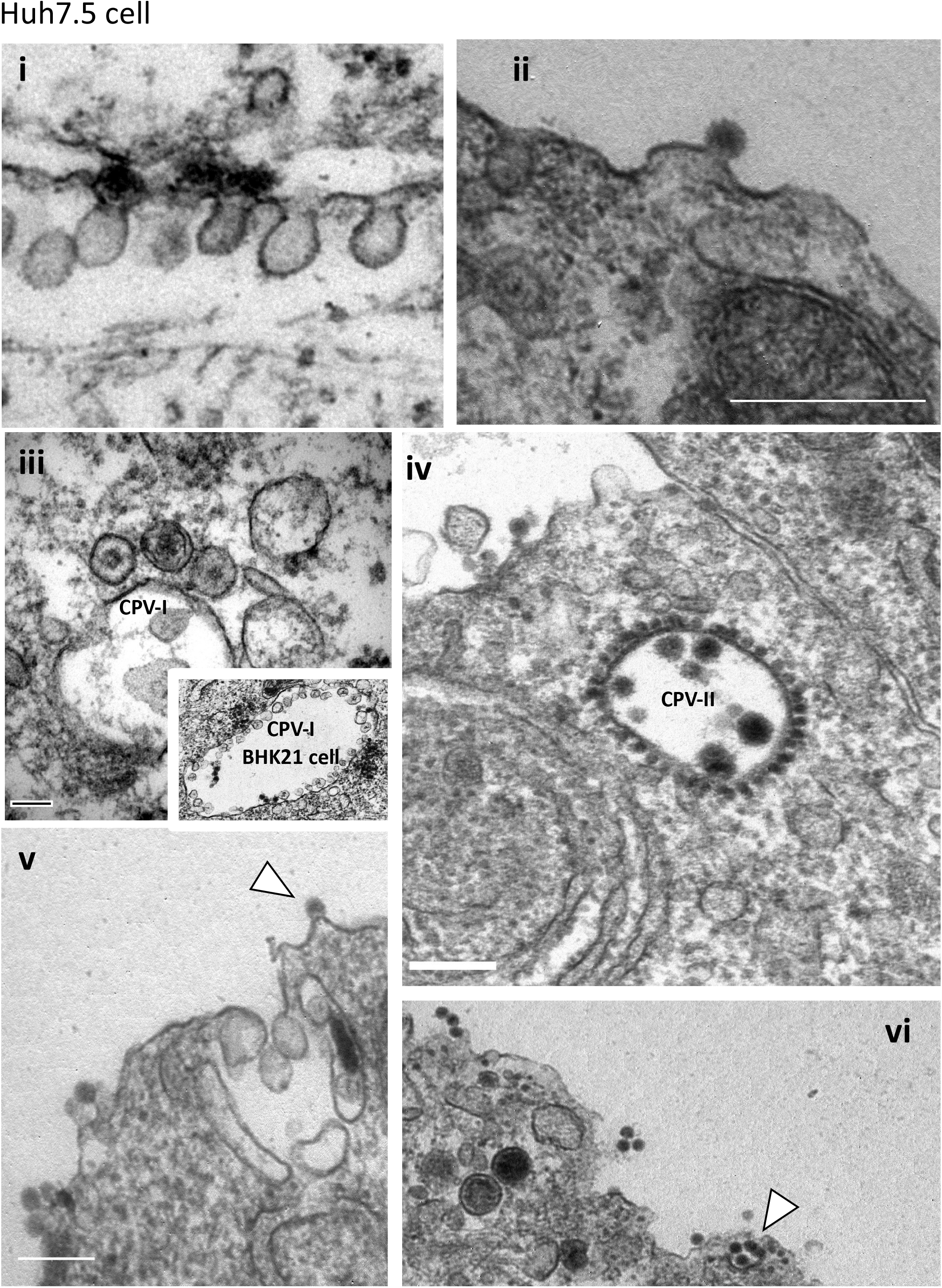
Electron microscopy analysis of MAYV lifecycle in vertebrate cells. A) Huh7.5 cells infected with MAYV at an MOI of 5, fixed and sectioned for TEM analysis. Virus entry by (i) clathrin mediated endocytosis pathway and (ii) direct membrane fusion. Cytoplasmic replication spherules with central electron dense structures (iii) and (iv) formation and maturation of nuclear core (NCs) in cytoplasm. v) and vi) release of viral particles by membrane budding and exocytosis. RS=Replication Spherules; CPV: Cytopathic Vacuoles.

In C6/36 cells, viral entry through RME was seen (Fig. 7i). Furthermore, similar to the infection of Huh7.5 cells, replication spherules (Fig. 3D), NC-containing vesicles, budding virus, and exocytic vesicles carrying mature viral particles were detected in the infected mosquito cells. In contrast to the series of membrane-attached spherules facing inside the CPV-I of BHK-21 cells, in C6/36 cells, most of the clustered spherules were unattached to the vesicular membrane (Fig. 7ii, iii). NCs were also observed inside and nearby the CVP-II membrane (Fig. 7iv), demonstrating that MAYV maturation and replication occur in the same cellular space. Intermediary CPV-I and CPV-II replication spherules and internally budded viral particles were observed in both early and late phases of infection (Fig. 7ii, v). These modified membrane structures were also seen near the rough endoplasmic reticulum and Golgi complexes. Furthermore, NCs close to the PM for budding and internally budded mature viral particles inside secretory intraluminal vesicles were seen (Fig. 7vi, vii).

**Figure 7.**
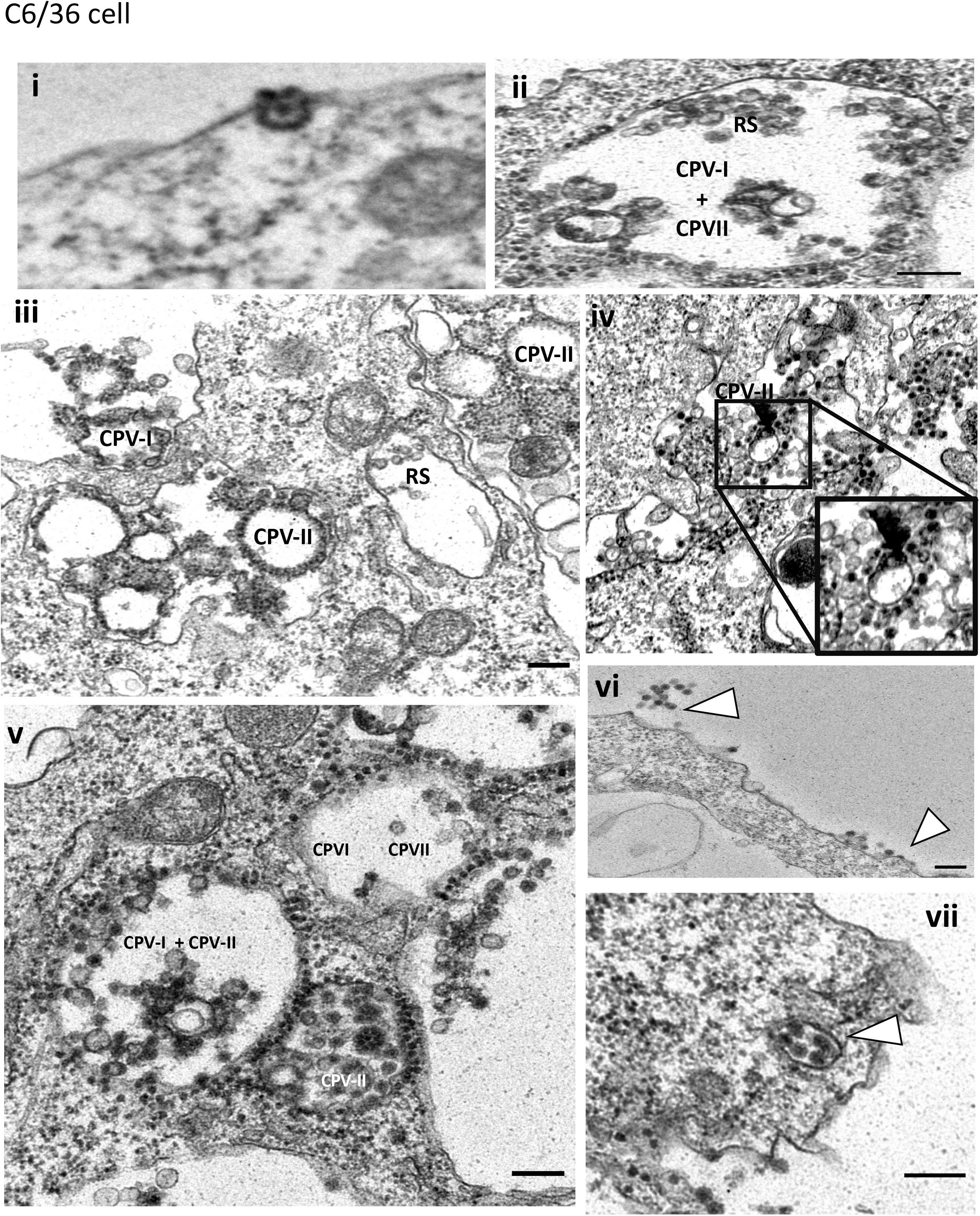
Electron microscopy analysis of MAYV lifecycle in invertebrate cells. C6/36 cells were infected with MAYV at an MOI of 5 and fixed for TEM analysis. i) Virus entry via clathrin coated pits like vesicle. ii) and iii) Virus replication inside the replication spherules in the cytopathic vesicle and on the cell membrane. ii) and vi) Virus replication and maturation in cytopathic vesicle I and II hybrid vesicles. v) formation and maturation of nuclear core (NCs) in cytoplasm. Release of viral particles by vii) membrane budding and viii) exocytosis. RS=Replication Spherules; CPV: Cytopathic Vacuoles

### MAYV induces both mitochondrial dependent and independent apoptosis

MAYV replication in Vero, BHK-21, and Huh-7.5 cells resulted in cytopathological changes characterized by cellular fusion and multinucleated giant syncytia, detachment of infected cells, and eventually cell lysis and death. To assess the involvement of MAYV in the apoptosis pathway (Huh7.5 cells), a cell viability test, and a PARP and Caspase detection study were performed.

Cell viability of MAYV infected Huh-7.5 cells was quantitatively measured through neutral red uptake assay. Neutral red is a eurhodin dye, actively transported into live cells, which stains lysosomes and is subsequently measured to determine cell viability. Cell viability was reduced up to ∼40% at 24hpi and more than 90% at 48hpi (Fig. 8A). A series of morphological changes, including cytoplasmic blebbing, mitochondrial swelling, chromatin condensation, and nuclear fragmentation, were observed in the ultrastructural analysis of MAYV infected cells (Fig. 8B).

**Figure 8.**
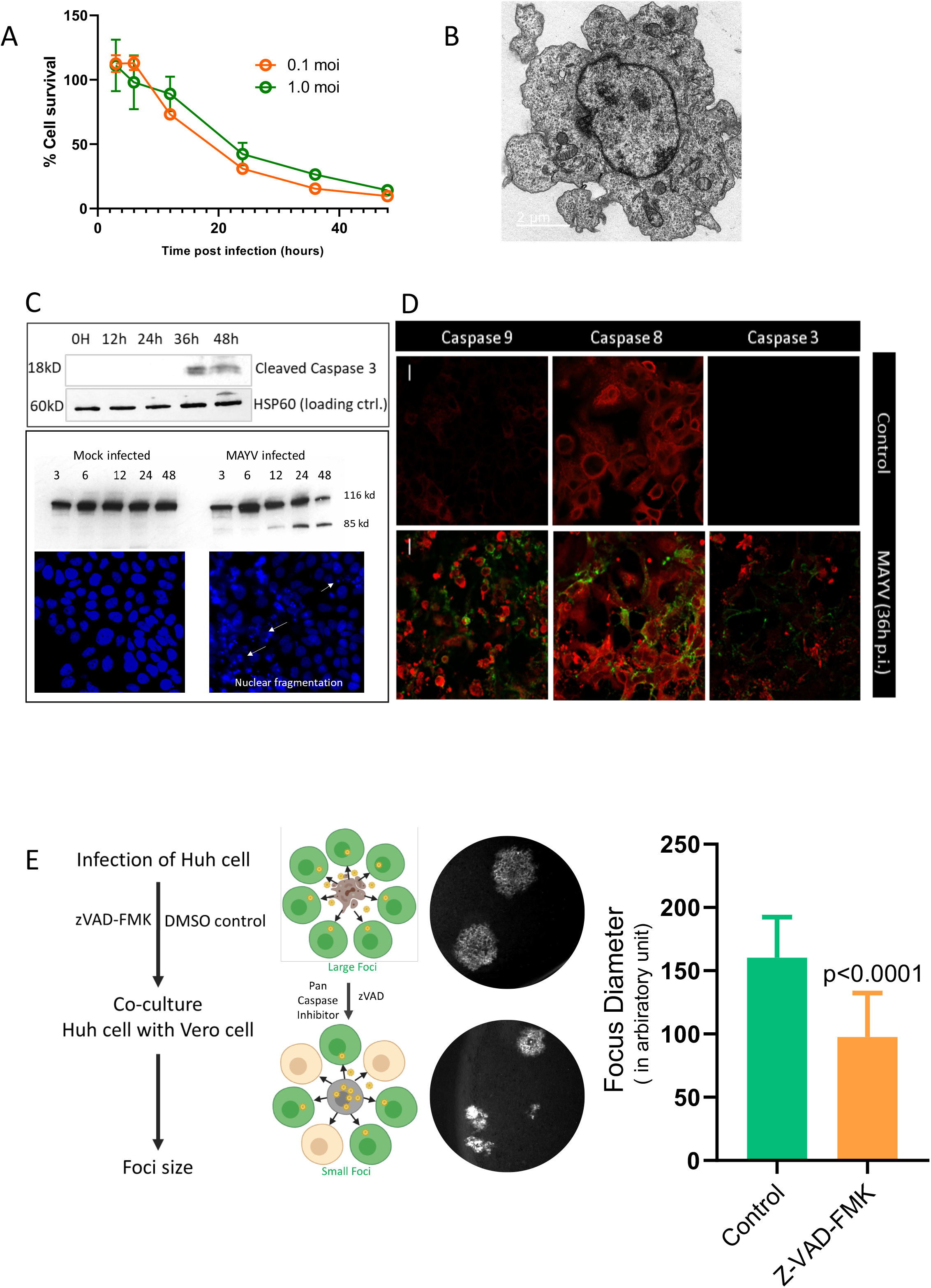
Both intrinsic and extrinsic apoptotic pathways are activated during MAYV infection. A) At 3, 6, 9, 12, 24, 36, and 48hpi Huh7.5 cell viability was measured by neutral red uptake assay. Data from three independent experiments were plotted, error bars indicate standard deviations of the means. B) TEM analysis showing apoptotic bleb and nuclear fragmentation of MAYV infected Huh7.5 cell. C) The cleavage of Caspase 3 at 12, 24 and 48hpi and PARP at 3,6,12, 24 and 48hpi by Western blot analysis in the cell lysate of mock and MAYV Huh7.5 cells. Uncropped blots are provided as Supplementary Figure 1. Hoechst 33342 staining of MAYV infected Huh7.5 cells (36hpi) observed under UV light. Arrows (bottom righthand side panel) indicate nuclei that contain condensed chromatin. D) Immunofluorescence detection of Cleaved Caspase 9 (intrinsic apoptotic marker), Caspase 8 (extrinsic apoptotic marker) at 36hpi and cleaved caspase 3 in MAYV infected Huh7.5 cells. E) Focus size in Vero cells by MAYV infected Huh7.5 cells treated with apoptosis inhibitor (zVAD-FMK) or vehicle control (DMSO).

Poly (ADP-ribose) polymerase (PARP) is an enzyme that suppresses nuclear fragmentation and apoptotic body formation. Under cellular stress such as viral infection, PARP is cleaved and compromises the cellular viability. Cleaved PARP, a marker of cells undergoing apoptosis, is detected at 12hpi and notably at its peak at 24hpi in the immunoblot of cells infected with MAYV. Additionally, stained cells with Hoechst 33342 revealed that MAYV induced chromatin condensation, nuclear fragmentation at 12hpi and is extensive around 36hpi, correlating with the PARP immunoblot results (Fig. 8C lower panel). Caspase-3, a cysteine protease, activates the PARP enzyme. Cleaved caspase-3 which is the active form of caspase-3 was detected at 36 and 48hpi (Fig. 8C upper panel) in the MAYV infected Huh7.5 cell lysates.

Caspase-9, an intrinsic pathway marker, and caspase-8, which initiates the extrinsic pathway, were detected in MAYV infected cells through immunostaining at 12hpi and 24hpi, respectively. Furthermore, active caspase-3, which cleaves the PARP enzyme and culminates in nuclear fragmentation and cell death, was detected at 12hpi in the immunoblot. This suggests that apoptosis which is led by the extrinsic pathway due to MAYV infection is subsequent to the early intrinsic pathway (Fig. 8D).

Finally, to test the hypothesis that MAYV utilizes the apoptosis pathway to maximize its spread, a co-culture experiment was designed. MAYV Huh7.5 infected cells pretreated with caspase inhibitor (ZVAD-FMK) were co-cultured with Vero cells that served as the infection center. The focus size formed by the caspase inhibitor treated cells was reduced up to 50% compared to the control group indicating that MAYV utilizes the apoptosis process to maximize the infection process (Fig. 8E).

## Discussion

Like other members of alphaviruses, the life cycle of MAYV is rapid and infectious virus can be detected in the culture supernatant as early as 3-5hpi in vertebrate cells and 7-9hpi in mosquito cells [13]. The viral titers are similar in mosquito and vertebrate cells at 24hpi and the viral titers in supernatants reach 10^7^–10^8^ FFU/ml within 48h, depending on the cell types. Further in agreement with previous reports of transmission of MAYV by *Anopheles* mosquitoes [8], Sua5b, a cell line derived from *Anopheles gambiae*, supported efficient replication and produced high viral titer similar to *Aedes albopictus* C6/36 cells. The lower viral titer in Aag2 cells compared to other insect cells and early plateau (36hpi) is not surprising, as Aag2 cells are persistently infected with insect-specific viruses which may possibly interfere with replication of MAYV [14]. MAYV had also been reported to replicate to high titer in avian cell lines which migratory birds has been hypothesized to be its reservoir [15].

Serological diagnosis is critical not only for disease surveillance but also for bedside diagnostics and is preferred over molecular assays especially for resource limited setup. The gold standard for serological diagnosis is the plaque reduction neutralization test, which demonstrates the virus-neutralizing capacity of serum samples [16]. Before any tests, the serum samples are heat-inactivated for 30 minutes at 56°C to inactivate complement proteins and any infectious viral particles, if there are any. Sometimes the inactivation time needs to be prolonged (e.g., 60 minutes for Western equine encephalitis virus and CHIKV), ensuring the sensitivity of the assays and the safety of laboratory personnel [17, 18]. We tested and compared the thermal stability of MAYV along with related Alphaviruses SINV and ONNV. After 3h of incubation at 45°C some of the virus particles were still infective but when the temperature increased to 55°C all of them lost their infectivity. The complete inactivation of MAYV could be achieved after 60 minutes of incubation at 55°C.

*In vitro* characterization of MAYV showed overt cytopathic effects in vertebrate cells, which facilitated the development of a plaque assay. The viral strain used in this study forms a mixture of large and small plaques with clear and sharp boundaries demonstrating the presence of a swarm of viral population with different viral entry and replication capacities. In contrast, mosquito cells did not show any cytopathic effect, were chronically infected with MAYV, and continuously shed infectious viral particles. This demonstrates that MAYV has developed a delicate balance between the mammalian host and mosquito host that allows its persistent survival in nature.

The role of RME has been established as one on the major route of entry for alphaviruses [10, 13]. As the endosome migrates closer to the nucleus, the inner compartment becomes acidic. This acidification causes the glycoproteins on the virus surface to undergo conformational changes and fuse with the endosomal membrane and releases the viral genome into the cytoplasm [19].

Mxra8, an adhesion molecule primarily expressed on epithelial myeloid, and mesenchymal cells, has been recently established as a receptor for CHIKV, ONNV, RRV, and MAYV [20]. Electron dense clathrin-coated pits in MAYV infected cells and significant reduction of viral titers both in the vertebrate and invertebrate cells upon chemical inhibition of dynamin-2, a key protein required for the formation of clathrin-coated pits and vesicles, suggests the role of RME in MAYV entry [21]. Further, the use of NH_4_Cl and Bafilomycin that alters the pH of the early and late endosomal vesicles inhibited the release of the viral genome into the cytoplasm and infection process in turn and production of infectious virus [22]. This indicates that the release of the MAYV genome is mediated through pH-dependent RME.

Upon releasing viral RNA into the host cytoplasm, which has the same polarity as that of cellular mRNA, it undergoes several rounds of translational events using host cellular machinery generating viral polyproteins. These are further processed by cellular and newly synthesized viral proteases creating the viral replication complex. The nonstructural proteins of alphaviruses induce intracellular membrane remodeling that results in the appearance of cytopathic vacuoles (CPVs) and has been greatly studied using the SFV model. The CPVs are of two types, CPV type I (CPV-I) and CPV type II (CPV-II) [23]. Double labeling of organelle and viral nonstructural proteins showed that CPV-I are derivatives of late endosomes and lysosomes, while monensin treatment results in accumulation of E1/E2 glycoproteins of SFV in the trans-Golgi network (TGN), indicating TGN origin of CPV-II [24, 25]. The replication complexes are strategically concentrated in CPV-I and replication spherule for efficient viral genome replication and to escape the cellular antiviral response [26]. These replication spherules that has access to the host cellular raw materials through an opening toward the cytoplasm were seen either inside CPV-I or transported to the plasma membrane [27]. Both vertebrate and invertebrate cells had similar spherules with a central electron-dense material, possibly the MAYV replicating RNA. dsRNAs are unique to viral infected cells and are the markers of replication intermediates/replication complex [28]. In the confocal image, detection of dsRNA in the plane with the nucleus indicated the replication of MAYV in the cytoplasmic CPV-I, which appears during the early stage of infection and above the plane of the nucleus, indicating the presence of replication complex on the cell membrane that is further validated in the electron micrographs. Like other alphaviruses, CPV-II was the predominant vacuolar structure in the later stage of MAYV infection both in Huh7.5 and C6/36 cells. Our electron micrograph analysis showed three main different populations of CPV-II in Huh7.5 cells. The first population consists of numerous nucleocapsids (NCs) attached to the cytoplasmic face of membranes, second has NCs enclosed inside the vesicles, and the third were transporting vesicles containing E/E2 viral glycoproteins from the TGN to the viral budding sites on the plasma membrane. In contrast to previous reports NCs were seen in the cytoplasm close to the plasma membrane of infected Huh7.5 and C6/36 cells. The assembly of NCs as matured viruses was observed to take place either by a budding process from the plasma membrane into the extracellular space or mature inside the CPVs transported through the exocytosis pathway. The C6/36 cells had additional hybrid vesicles containing both replication spherules and NCs, also having vesicles with mature virus particles.

Arthritogenic alphaviruses, including MAYV, have been associated with inflammation of the joints and often with other tissues. Inflammation and apoptosis go in parallel with disease severity. Although apoptosis eliminates infected cells, viruses have evolved to manipulate this cellular antiviral mechanism for maximizing the production of progeny virus and their spread to maintain a prolonged and high viremia in the vertebrate host, a property that is relevant especially for arboviruses for their maintenance in the natural transmission cycle. CHIKV and SINV camouflage in apoptotic blebs to facilitate infection of neighboring cells. Apoptotic blebs bodies are quite evident in the MAYV infected cells. Most commonly, blebs are seen during apoptosis and contain part of the cytoplasm (∼2%) with or without organellar fragments for recycling by the phagocytic cells [33, 34]. We designed a coculture-infection center assay that produced a smaller diameter focus size in the presence of apoptosis inhibitor ZVAD-FMK that inhibits apoptosis at an early stage, indicating the role of apoptosis in the spread of MAYV infection. Up-regulation of proapoptotic proteins or down-regulation of antiapoptotic proteins can alter mitochondrial membrane permeability that can promote the release of cytochrome c [29]. Cytochrome c interacts with Apaf-1 and caspase-9 and forms a multiprotein complex apoptosome, leading to activation of caspase-3 and then apoptosis [30,31]. Increased activation of caspase 9 during MAYV infection indicates the involvement of mitochondrial intrinsic pathway. Further detection of active caspase 8, possibly activated by secretion of cellular or virus-induced death signal by cell membrane death receptors, demonstrate that the MAYV induced apoptosis is triggered through both intrinsic and extrinsic pathways.

In conclusion, we present a comprehensive *in vitro* characterization of MAYV in both vertebrate host and invertebrate mosquito vector cells. Besides pH-dependent receptor-mediated endocytosis (RME) MAYV uses macropinocytosis for entry into vertebrate cells. During acute infection in vertebrate cells, MAYV achieves very high titer and utilizes the apoptotic pathway for efficient transmission to neighboring cells. In mosquito cells, a burst in high viral titer follows continuous virus production at reduced levels, indicating the evolutionary adaptation of arboviruses with their mosquito vector. Further, it will be interesting to understand and identify the molecular signature(s) that makes them clinically different from their close geographical ancestors.

## Acknowledgements

We thank Missy Hazen for assistance with microscopy. This work was funded by NIH grants R01AI150251, R01AI128201, R01AI116636, and R21AI128918, USDA Hatch funds (Accession #1010032; Project #PEN04608), and a grant with the Pennsylvania Department of Health using Tobacco Settlement Funds to JLR, and grant NIH R21AI128918, a seed grant from The Penn State Huck Institutes of the Life Sciences, and startup grant and ASPIRE grant from the University of South Carolina to SP.

## Authors Contributions

Conceptualization: SP, JLR; Methodology: SP; Data Curation: SP, MB, CCH, RS, ML; Funding acquisition: JLR, SP; Writing-first draft: SP; Writing-review and editing: SP, JLR, SH, MB, CCH, RS and ML; Writing-finalizing MS: JLR and SP.

## Notes

### Competing Interest Statement

The authors have declared no competing interest.

### Summary of Updates

Manuscript has been revised to reflect new experimental results and an additional ooauthor

