## Supplementary Figure 1 for "Everything you wanted to know about Mayaro virus but were afraid to ask: Characterization and lifecycle of Mayaro virus in vertebrate and invertebrate cellular backgrounds"

Sup Fig 1

Mock infected MAYV infected

3 6 12 24 48 3 6 12 24 48

Cleaved PARP

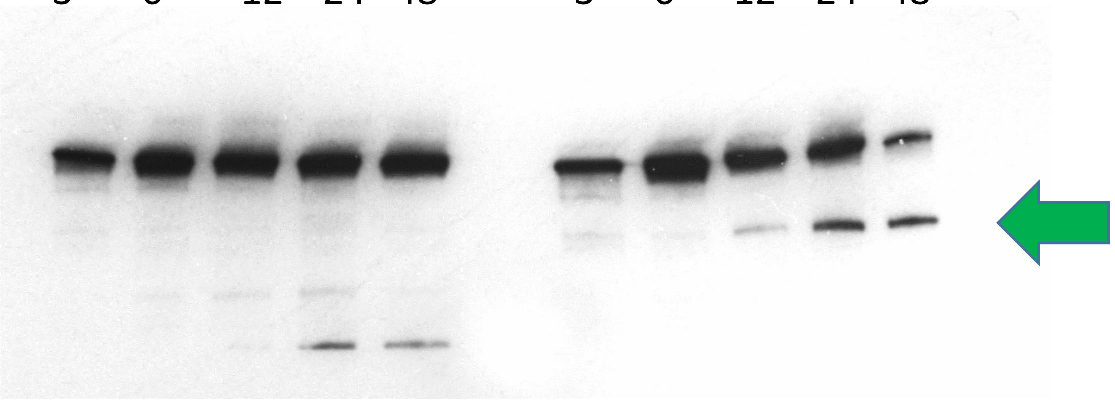

Cleaved Caspase3

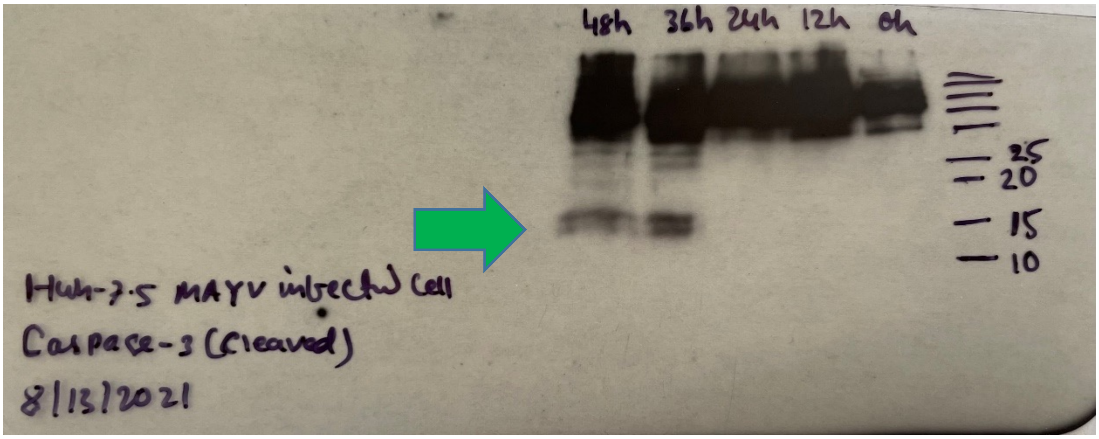


HSP60 Loading control for Cleaved Caspase3

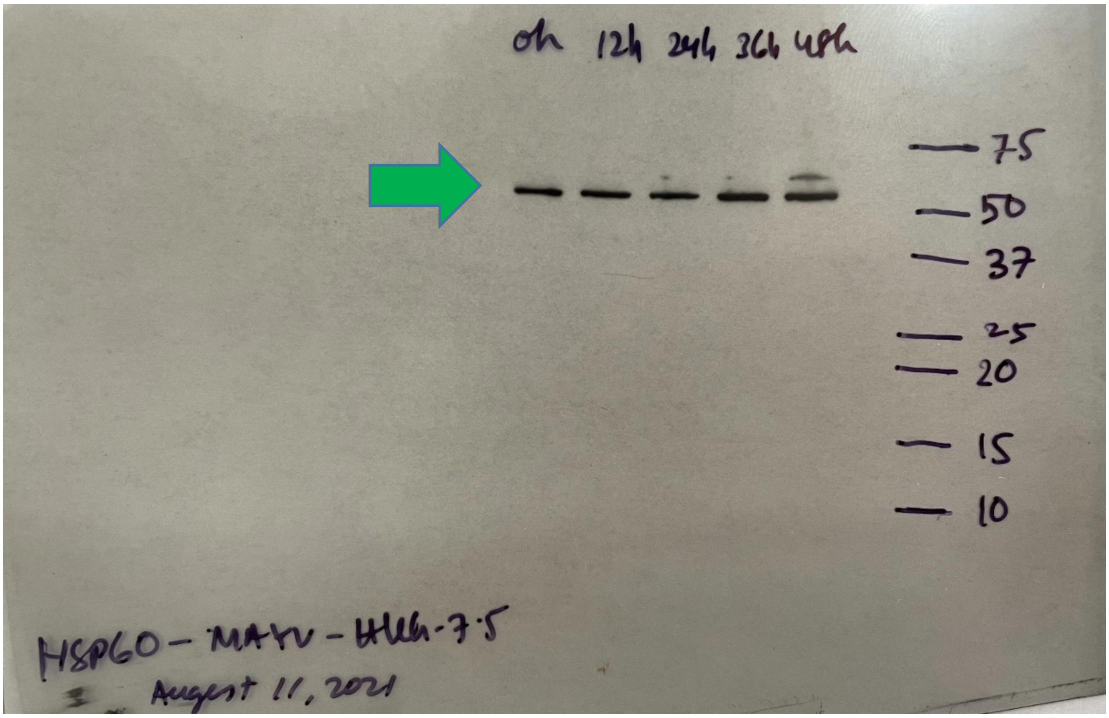
